# A Machine Learning Application to Camera-Traps: Robust Species Interactions Datasets for Analysis of Mutualistic Networks

**DOI:** 10.1101/2025.03.01.640990

**Authors:** Pablo Villalva, Pedro Jordano

## Abstract

Recording and quantifying ecological interactions is vital for understanding biodiversity, ecosystem stability, and resilience. Camera traps have become a key tool for documenting plant-animal interactions, especially when combined with computer vision (CV) technology to handle large datasets. However, creating comprehensive ecological interaction databases remains challenging due to labor-intensive processes and a lack of standardization. While CV aids in data processing, it has limitations, including information loss, which can impact subsequent analyses.

This study presents a detailed methodology to streamline the creation of robust ecological interaction databases using CV-enhanced tools. It highlights potential pitfalls in applying CV models across different contexts, particularly for specific plant and animal species. The approach aligns with existing camera trap standards and incorporates complex network analysis tools. It also addresses a gap in ecological research by extending the methodology to behavioral studies using video-based image recognition, as most current studies rely on still images.

The study evaluates CV’s performance in estimating species interaction frequency (PIE) and its ecological implications, with examples of plant-frugivores interactions for seed dispersal. Results show that up to 10% of pairwise interactions may be missed with CV, with information loss varying among focal species and individual plants. This poses challenges for individual-based approaches, where unbiased data collection requires extra caution. However, the loss is minimal compared to the vast data CV enables researchers to gather. For community-level approaches, only three out of 344 unique pairwise interactions were missed, and overall estimates of both PIEs and interaction strengths remained largely unaffected.

The methodology provides a valuable resource for ecologists seeking to document ecological interactions efficiently. It offers guidelines for collecting reliable data while addressing CV’s limitations in capturing unbiased species interaction data. Despite its constraints, CV significantly enhances the ability to gather large-scale interaction data, particularly at the community level, making it an indispensable tool for ecological research.

## 1. INTRODUCTION

No species on Earth lives without interacting with other species, the reason why ecological interactions are at the core of the Web of Life (Thompson 2009). These interactions play a vital role in supporting Earth’s systems and are crucial for understanding ecosystem functioning (Loreau, 2001), leading to an increasing consideration as part of biodiversity (Jordano 2016a, Pollock et al., 2020). To better comprehend the contribution of biodiversity to ecosystem functioning, there is an urgent need to record, quantify, and assess ecological interactions.

Frugivory, an essential plant-animal interaction, has been shaping the co-evolution of animals and plants for over 80 million years (Tiffney 2004, Erikson, 2014). Animals benefit from pulp-derived resources provided by plant fruits, while plants rely on frugivorous animals for seed dispersal services, making seed dispersal a central process in the natural regeneration of plant populations (Jordano, 2014). However, monitoring frugivory presents a significant challenge due to its labor-intensive nature, often resulting in incomplete data samples, a problem analogous to any biodiversity sampling (Gotelli and Colwell, 2011, Jordano 2016b). Moreover, comparing or replicating results from different data sources (e.g., interaction records derived from direct watches, DNA barcoding, camera traps, etc.) poses limitations (Quintero et al., 2022), yet ultimately results in more complete samplings. Fortunately, recent technological advances have enabled the acquisition of high-quality field data and sufficient computational power to process large datasets. This progress provides an opportunity to document frugivore assemblages in-depth, moving beyond the community scale and enabling the assessment of processes at finer scales such as recording interactions for individual plants to analyze ecological correlates of interaction patterns within populations (e.g., Miguel et al. 2018).

Over the last two decades, the use of camera traps for wildlife monitoring has significantly advanced our understanding of vertebrate ecology and population structure (O’Connell et al., 2011). It is a cost-effective method for monitoring multiple species over large spatial and temporal scales, however the time required to process the data can limit the efficiency of broad scale surveys. These remote cameras have been employed in various studies, ranging from population (Bischof et al., 2020; Gardner et al., 2010) and community-scale (Ahumada et al., 2011; Wittische et al., 2021) investigations to species distributions (Rich et al., 2017; Tobler et al., 2015). Camera traps are a non-invasive method with wide application in behavioral studies (Caravaggi et al., 2017, 2020) as it offers information that allows to record natural behaviors of animals without human disturbance in natural environments. Several studies have been focusing on diverse behavioral aspects, including activity periods (Suselbeek et al., 2014), daily activity patterns (Frey et al., 2017; Leuchtenberger et al., 2014; Ferreiro-Arias et al. 2021), road crossing behavior (Villalva et al., 2013), human-wildlife conflicts (Johnson et al., 2006), scent marking behavior (Rafiq et al., 2020), among others. However, despite their usefulness, complex ecological processes such as species interactions have only been inferred using camera traps (Beirne et al., 2021; Clare et al., 2016; Niedballa et al., 2019; Selwyn et al. 2020), and the application of camera traps in behavioral studies, particularly regarding foraging behavior (see, e.g., Koike et al. 2012) is still in its infancy (Caravaggi et al., 2017).

The use of video recordings focusing on specific plants allows for the documentation of detailed interaction data, including behavioral information, concerning interacting species. However, effective pipelines for handling large volumes of videos currently lack standardization and are underdeveloped for ecological purposes. Fortunately, recent advancements in Artificial Intelligence (AI) applied to image recognition (Leorna and Brinkman, 2022; Rigoudy et al., 2022; Velez et al., 2023; Petroni et al. 2024), along with the decreasing cost of camera trap devices, have made it feasible to collect and manage high-quality data within a reasonable timeframe and with an affordable budget. AI enables the processing of millions of images in a short time. Though, substantial and diverse pre-processed data is necessary to train models, including pre-processing of images, model training, classification, manual quality checks, and data formatting (Bohner 2023, Celis et al. 2024). Although AI has demonstrated accurate species identification (Carl et al., 2020; Gómez Villa et al., 2017; Whytock et al., 2021) and counts (Norouzzadeh et al., 2018), current models have primarily been trained for mammals due to the greater availability of information for them (Burton et al., 2015). In contrast, bird species image recognition remains limited (but see Cornell Lab for image and sound recordings). Furthermore, the performance of AI approaches might be compromised when models are developed using training datasets that are unbalanced (Gomez Villa et al., 2017) or small and geographically limited (Schneider et al., 2020; Tabak et al., 2019), or when applied to low-resolution images (Gómez et al., 2016). Additionally, the process of model creation and refinement demands technical and programming expertise that extends beyond the capabilities of many ecologists (Christin et al., 2019; Tabak et al., 2020). Nevertheless, current AI models can accurately classify images to remove blank (empty) ones (Berry et al., 2019) making it possible to select for detailed review of just those images containing animals, notably reducing the revision effort while minimizing the tagging error by low-trained models. Our protocol takes a conservative approach and does not aim to automatically identify species, as we recognize that identification at the species level still requires extensive model training and obtaining fine grain behavioral information is still not possible. Instead, we leverage current image recognition techniques to eliminate blank images, with direct application to the estimation of species’ interaction frequencies (Villalva et al. 2024).

In this paper, we introduce a streamlined workflow for effectively managing a large volume of videos to gather data on plant-animal interactions. This workflow represents, to our knowledge, the first application of image recognition for ecological camera trap surveys specifically focused on videos rather than still images, and it establishes standardized procedures for the specific analysis of plant-animal interactions data. Our primary goal is to reduce effort and time in data compilation by offering a method that is suitable for multi-species monitoring and adaptable to various ecosystems. The approach integrates a field protocol based on camera trap operation and data standards with available advanced AI for image recognition, complemented by a viewer program to optimize image review and tagging. We also offer several approaches to minimize time consuming data management such as automating the acquisition of sampling effort data and inference of actual interactions from video sequences of short duration. The main objective is to construct an accurate and fully annotated dataset for documenting the presence and frequency of plant-animal interactions records while time and effort required is minimized.

## 2. METHODS

Our approach for monitoring plant-animal interactions in natural habitats involves the strategic placement of camera traps, aimed at specific plant species, referred to as focal plants. We use replicated individual plants of the same species and deploy cameras in each one; results of such monitoring can thus be used as pooled, aggregated data for the species or analyzed to assess among-plant differences. The goal is to obtain accurate records of visits to the plants by animals allowing estimates of interaction probability (PIE, probability of interspecific encounter; Poisot et al. 2016). *PIE_ij_* is thus the probability that species *i* establishes an interaction with species *j*, or the probability that an individual plant, *i*, interacts with animal frugivore species *j*. These cameras are set in video mode, an added value to provide valuable insights for species identification and behavior, as well as an accurate quantification of fruit consumption and fruit handling, which become possible at least in some of the sequences obtained.

Interaction frequencies, *PIEs*, etc. can be derived by tallying those video sequences with evidence of an animal frugivore present while foraging at the focal plant. Active foraging for fruits by the animal can be evidenced by the video sequence showing an actual event of fruit picking and, maybe, fruit ingestion or fruit dropping. The interaction can also be inferred when the animal is visualized on the plant when actively foraging (despite no evidence of fruit feeding); and in order to not overestimate interaction frequency, such foraging event can be weighted by the actual proportion of fruit food in the diet of the animal species recorded (Quintero et al. 2024). Due to the typically short recording times of the cameras when triggered (usually 10-20s sequences), interaction frequencies can be drastically underestimated if just those sequences with actual fruit handling/ingestion are considered: a high frequency of false negatives would be ignored, when the short recorded sequence is actually part of a feeding sequence at the plant including fruit ingestion. It is extremely rare that video sequences of 10-20s include the total length of a frugivore’s visit to a fruiting plant.

Video-based monitoring thus offers several advantages over traditional camera-trap techniques that rely on still pictures. By capturing movement, we are able to identify animal species and their behavior, thereby gaining a deeper understanding of the interaction event. In addition, for some video recordings we can accurately measure fruit consumption and determine fruit feeding behavior and consumption rates (see, e.g., Snow and Snow 1988, Levey 1987, Jordano and Schupp 2000, Moermond and Denslow 1985 for a full description of such diversity of fruit feeding behaviors). However, video-based monitoring generates a significant demand for high digital storage compared to still images, which must be considered. Through video recordings, we gain insights into the importance of different plant species for wildlife within the ecosystem, and the ecosystem services provided by animals for plant dispersal.

### Image recognition model

Camera-trap monitoring presents challenges, especially when using a large number of devices in windy environments. Under wind conditions, the movement of grasses and tree branches can easily lead to incorrect triggering, resulting in a significant number of empty images. This issue becomes particularly relevant in plant-animal interaction surveys focused on frugivory, where the natural environment must be unaltered and removal by hand of grass or branches is not an option. Thus empty images may constitute a substantial portion of the video collection. Consequently, reviewing all recorded videos to identify ecological interactions becomes unfeasible. To address this challenge, several image recognition models have been developed to eliminate empty images (Chalmers et al. 2023, Ahumada et al. 2020, Tabak et al. 2022). In our pipeline, we used Megadetactor, a ready to use image recognition model that has been trained in diverse ecosystems worldwide and demonstrated high performance, effectively removing blank images from the image data (Beery et al., 2018).

### Protocol pipeline

Our streamlined workflow comprises three essential stages: (1) pre-processing, (2) video processing, and (3) post-processing (Fig. 1). During stage-1, we establish a standardized data structure and procedure for camera trap settings and sampling effort automatization. This ensures the consistency and accuracy of data collection avoiding early errors that would cascade down the process chain. In stage-2, we leverage image recognition to classify and select just the videos featuring animals for further visualization. Additionally, we use time-saving software to visualize and tag the selected videos, optimizing the extraction of valuable information. Finally, in stage-3, we integrate datasets from different video batches, encompassing various seasons, focal species, or camera sources, and consolidate them for comprehensive analysis. This final step results in a cohesive dataset, providing a solid foundation for further exploration through network analysis tools or other methods for interpretation.

**Figure 1.**
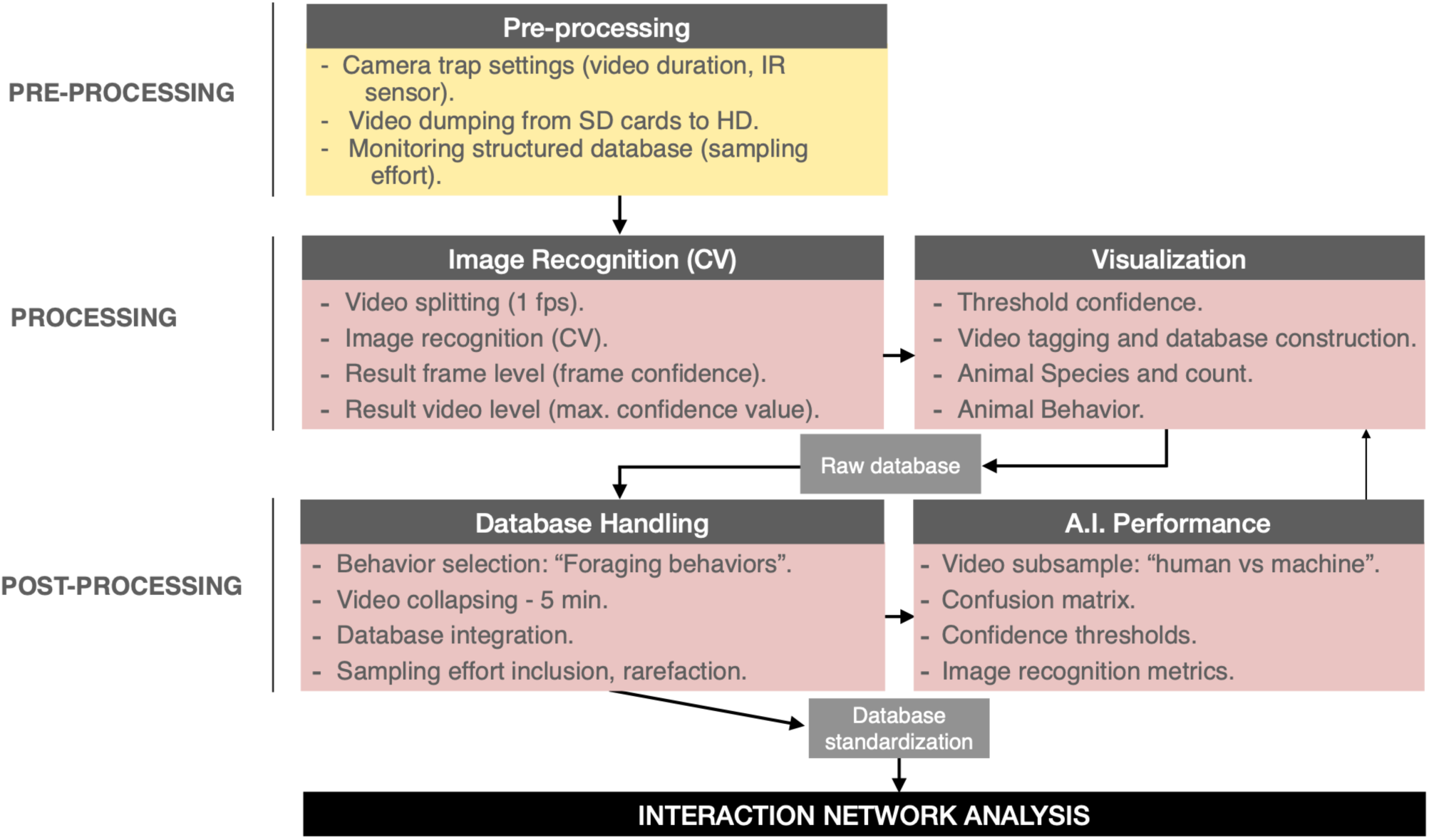
Detailed workflow for optimizing data obtained during sampling of interaction events between animal frugivores and fleshy-fruit producing plant species using camera traps deployed in the vicinity of fruiting plants. The raw data obtained represent the interaction frequency between each animal species and the plant species visited, estimated from the number of 10 s video sequences obtained during camera operation whenever the camera is activated by movement by the frugivore species. The protocol includes steps for data pre-processing, splitting of sequences to avoid pseudo-replicated records, proper identification of sequences containing animal activity (i.e., the camera was actually triggered by animal activity and not by wind, etc.), and sequence analysis and compilation of distinct interaction records.

**Figure 2.**
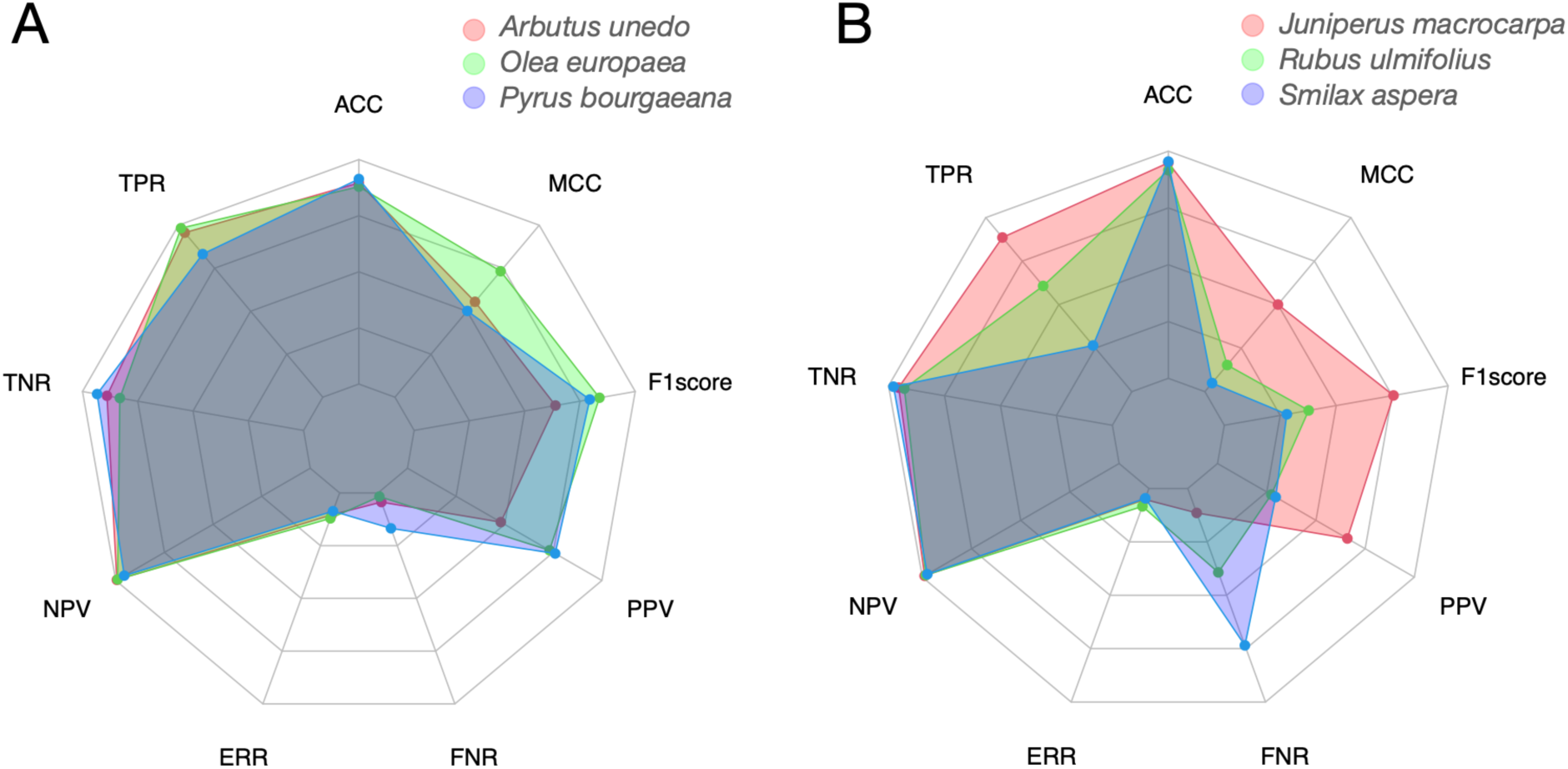
Performance metrics for six distinct plant species, represented by different colors in spider web plots, evaluated for nine different metrics (radii in the plots). On the left panel, three species exhibit consistent and satisfactory performance metrics, whereas the right panel illustrates three species with varying performance, including two with poor results. Performance metrics used: [TPR] True Positive Rate (Sensitivity, Recall), [TNR] True Negative Rate (Specificity), [FPR] False Positive Rate (Fall Out), [FNR] False Negative Rate (Miss Rate), [PPV] Positive Predictive Value (Precision), [NPV] Negative Predictive Value, [FDR] False Discovery Rate, [FOR] False Omission Rate, [ACC] Accuracy, [ERR] Error Rate, [F1score] F1 Score (Harmonic mean between TPR and PPV), [MCC] Matthews Correlation Coefficient.

**Figure 3.**
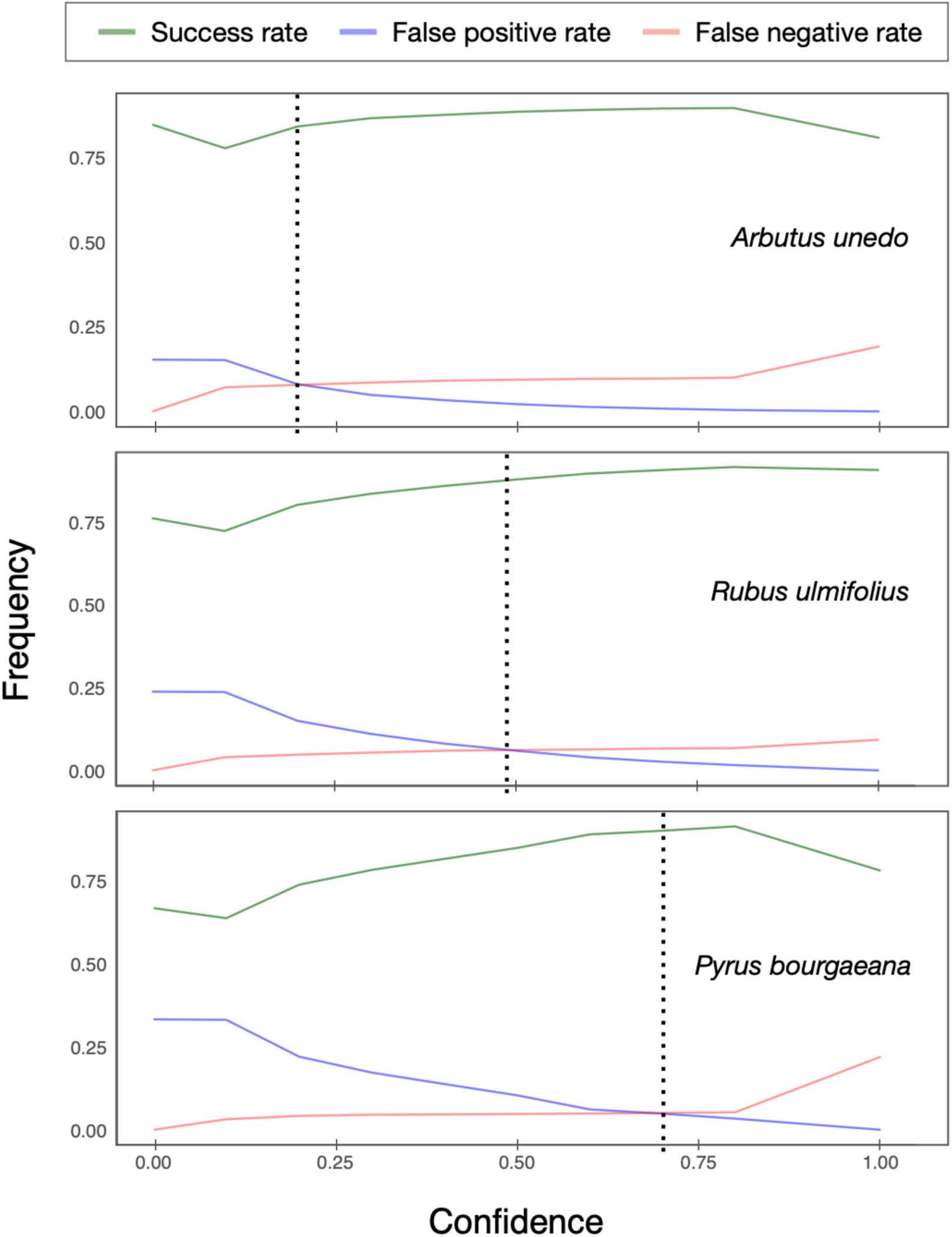
Cumulative curves depicting success (green line), false positive (red) and false negative (blue) rates relative to confidence level for three plant species, highlighting potential confidence thresholds. A dotted vertical line indicates the species-specific confidence threshold, where false positives and false negatives are minimized, while ensuring a high success ratio.

### a. Stage-1. Pre-processing

The deployment of multiple cameras in replicated positions requires regular checks, resulting in a large number of videos with identical names and dates. Thus, it is essential an effective organization and management of such a vast and intricate dataset for a successful database creation. The database structure needs to facilitate tracking the data at every stage of the process. To address this requirement, we adapted the CamTrap Data Package structure (Bubniki et al., 2023) and developed a standardized framework for controlling camera-trap data for frugivory-interactions data gathering. This structure ensures consistency and minimizes the risk of errors. It comprises three related plain texts containing all essential information for convenient data management and analysis (Table 1). For more detailed information and templates related to our customized structure refer to the supplementary material S1.

**Table 1.** The main data is structured in three related plain text files (.csv). Table *deployment* includes deploymentID, Location and camera Setup information for each camera. Table *video* contains information related to each field revision including dates and TimestampIssues. Table *observation* contains video-level information obtained from visualization.

| File | Description |
| --- | --- |
| deployment.csv | Table with camera trap deployments. |
| video.csv | Table with media files captured by camera traps. |
| observation.csv | Table with observations based on media files (after visualization). |

Camera trap settings for frugivory data gathering involve several key factors. First, it is advisable to use a consistent camera trap model throughout the study to maintain data integrity and reduce bias since detection may vary among different camera models and makes (Palencia et al., 2021). If using multiple camera models is unavoidable, a systematic rotation of deployment could be considered, although this may complicate keeping track of camera identification through the workflow. The sensitivity of the camera trap should be set carefully, considering both the surrounding environment and the specific plant species under study. In open areas with minimal movement interference, a higher sensitivity can be set, while in more cluttered areas with higher movement levels, a lower sensitivity setting is recommended.

For optimal results, the camera traps should be set at the shoulder height of the target species, as this increases the probability of detection (Palencia et al., 2021). However, in a multi-species approach, achieving the ideal camera height placement can be challenging. To address this, we suggest using multiple camera deployments to capture a full range of visitors (e.g., ground vs. canopy). In these cases the field-of-view of each camera should not overlap to avoid duplication of data. A challenging aspect, almost with any plant species fruiting, is to optimally adjust the field of view while not sacrificing image detail with a deployment at a longer distance. Ideally, individual fruits should be visible in the image of the canopy, under good light conditions, but also with an adequate coverage of the maximum canopy area possible.

As stated above, video mode is necessary to accurately determine animal behaviors (and sometimes species identification). We recommend setting the video length consistently to 10-20s to conserve memory cards and batteries and streamline the post-process workflow. The motion trigger should have no more than 1s delay to maximize the recording of animal behavior in the scene. The longer length of 20s, or even longer, might be adequate if obtaining data on fruit foraging and feeding behavior is a relevant objective of the sampling, so that a longer part of the feeding sequence or even the full visit to the plant by the foraging animal can be captured. Systematically recording a file at the beginning and end of each deployment revision ensures an accurate control of the sampling effort. We offer a code snippet that automates the collection of sampling effort for each camera deployment (see *code* section).

### b. Stage-2. Processing

Automated processing of large video volumes poses significant challenges and time constraints, making their manual review costly and impractical. Artificial Intelligence (AI) for Image recognition technology (computer vision, CV) offers a promising solution by automating data collection. AI has demonstrated effectiveness in species recognition and identification, with studies reporting high accuracy rates. For instance, Norouzzadeh et al. (2018) achieved over 99% accuracy, Schneider et al. (2019) reported 93%, and Tabak et al. (2019) reached 97.6% accuracy using recognition algorithms. Yet, automated species identification needs to be further improved (Kissling et al. 2024) and it is essential to be cautious when interpreting model performance, especially when the model training dataset lacks diversity, as this restricts the generalization of the findings to other systems (Greenberg et al. 2019). Attempting to achieve similar accuracies as those provided above may prove unrealistic particularly when a diverse array of new species need to be identified in an untrained system, as is the case in most plant-animal interaction studies. Securing sufficient funding for the proper training of models to automate species identification might surpass the scope of available conservation funding for such projects.

#### Eliminating empty videos

A simpler recognition result can lead to significant gains in efficiently analyzing images. Using an object detector model to differentiate between empty and non-empty videos can save valuable time and is essential for streamlining the analysis process. Most field recordings from plant-animal interaction surveys contain unintended motion triggers caused by the movement of vegetation and thus do not contain relevant information. Removing these blank recordings can considerably reduce the review time. We employ MegaDetector (MD; Beery et al. 2019), an advanced object detection model capable of identifying animals, people, and vehicles in images. Although MD was originally designed for still images, our protocol represents a first attempt to apply object recognition to vast sets of camera trap video recordings.

#### Video splitting

The implementation of the object detector to video recordings requires splitting the video files into individual frames. To automate video splitting in large batches, we use an R script (see *code* section) that automatically extracts frames from video files and stores them in separate directories maintaining the parent file structure. Although video splitting is time-consuming, it offers a substantial advantage by significantly reducing the overall file storage size, particularly useful when uploading to an external server, a requirement for the use of GPU capabilities.

#### Applying the object detection model

After the video set is splitted up into individual frames, the model is executed on each individual frame. There are two primary methods to execute the MD object detection model:

1. Running the model on a local computer: The model can be freely obtained from the MD GitHub repository (https://github.com/agentmorris/MegaDetector). The convenience of running the model locally depends on the number of images to be processed, the computer’s hardware specifications, and the user’s proficiency in the Python programming language.
2. Submitting images to MD staff for model execution: This option is ideal for high-volume users who require access to high-performance processors. Users can submit images either through a physical hard drive or by uploading them to the cloud using the sftp protocol. It is essential to note that efficient recognition often requires high-performance computers due to the computational demands of the process. Additionally, whether the model is run locally or by the MD staff, it is advisable to perform a preliminary check by running the model on a few thousand images to ensure the effectiveness on the target dataset.

#### Model outputs

Once the model is executed a json file is generated as the main output. This standard data interchange format preserves the inherent structure of the input data, retaining the record of each processed frame file through the frame-level analysis. This frame-level output includes the probability, or confidence score, at which the model detects the presence of an animal (also person and vehicle, which are discarded for our specific purposes) in each frame. To obtain the video-level output, we aggregate the frame-level results for each video file by selecting the highest confidence value from the set of frames. This highest confidence value corresponds to the model’s confidence in detecting the presence of an animal in the video.

#### Visualization and database creation

Time-lapse software (Greenberg et al. 2020), an open-source tool designed for reviewing camera trap images and videos, proves to be an optimal solution for creating ecological-interaction datasets. It boasts good support for videos and a multitude of interface tools for accelerating the visual analysis and encoding. Time-lapse automatically extracts file information and metadata, presenting a custom interface for easy data entry and supporting visual searches. Time-lapse enables direct import of json outputs from the image recognition pipeline. This functionality allows data filtering based on the confidence level of the object detection model, which allows to implement the confidence threshold for excluding empty videos for visualization, streamlining the workflow.

The program allows the design of ad-hoc templates, which should match the structure of the “Observations.csv” file (see Supplementary material for detailed information). These templates are designed to ease the process of tagging information related to each video, such as species name, species count, behavior, etc. Ultimately, the output from Time-lapse can be exported as a plain text (.csv) file containing a list of videos with animals and their associated behaviors, along with relevant metadata information.

### c. Stage-3. Post-processing

#### Database handling

Each entry of the dataset is referred to an individual recording, which captures a 10-second duration video showing one or more animals exhibiting diverse behaviors, entailing the most detailed resolution of the dataset. Different behaviors may be selected for several behavioral studies. In our frugivory context, we focus on behaviors related to fruit consumption. Opting for a conservative approach, one could choose videos where the animals are observed directly handling and/or ingesting the fruit. However, this selection is likely to be overly strict, leading to an underestimation of the actual frequency of fruit feeding and thus of the interaction frequency, i.e., discarding as false negatives some records that, while not showing the animal handling the fruit, the foraging sequence captured is clearly part of a fruit-feeding bout an thus indicative of an actual interaction. Avoiding such potential underestimation of interaction frequencies involves including videos where the animals are likely within a feeding bout or clearly searching for food during the visitation sequence. This could encompass cases where the animal’s feeding action was not recorded due to its position or possibly occurred out of the camera’s view, probably due to the short duration (10s) of the recording. By incorporating these types of behaviors, a more comprehensive characterization of the actual interaction frequencies of these plant-animal interactions from the field survey can be achieved.

#### Video collapsing (temporal autocorrelation)

If the dataset remains unaltered after the selection of behaviors it will contain a significant bias due to temporal autocorrelation of consecutive 10s videos. To avoid introducing bias to the analysis, it is crucial to establish a baseline for defining unique frugivory interaction events. An objective criterion for defining events is to set a specific duration threshold to create independent events. In this protocol, we propose creating independent events of 5 min length. The rationale behind this approach is that a frugivore visit sequence starts when the animal arrives at the plant and concludes when it departs. These visits typically extend beyond a 10s length, during which the animal may or may not feed on fruits. Only in some instances the camera will record fruit handling and/or ingestion. And, probably, even in more rare instances, will record the whole visit duration. Pooling successive videos within 5-min intervals serves as an adequate way to represent a single interaction bout. Since events tend to be short, setting a 5 min duration will usually suffice, while allowing sufficient differentiation between visits from different individuals. This approach prevents the aggregation of clearly distinct visits of other individuals into a single interaction event.

#### Sampling effort inclusion

Sampling effort recording is crucial for controlling the level of effort for the different deployments, plant individuals and plant species, allowing to analyze and compare the amount of time spent on each of them, in order to evaluate the data collection process. In the final step of the post processing stage we include the information regarding sampling effort to the dataset. This information is automatically extracted from the parent directories at the video level by an ad-hoc code. The code relies on the metadata from the video files from a camera recorded during each revision, that are stored with this structure. As previously mentioned, triggering a first and last recording as explained in the pre-processing section is mandatory to ensure accurate sampling effort information. The code calculates the duration of each deployment based on the date and time of the first and last video, which represents the sampling effort for a specific deployment. The duration, expressed in decimal numbers (days), can be aggregated for individual and species-level. An overview of the code’s functionality is provided in the code section below and is available in the repository.

### d. Code

A brief definition for the code functions provided in our streamline protocol is provided below. The code is available in the Github repository: (https://github.com/PJordano-Lab/Ecological-interactions-camtrap-protocol).

- *Video splitting* - Automates the process of recursive extraction of frames from video files. It organizes the frames into new directories following the same structure as the original ones. The extracted frames are used for running the image detection model at frame level.
- *Working with json* - Processes JSON data, filters and selects specific video files based on their confidence levels, moves the selected video files to new directories with modified paths, and combines data from multiple JSON files into a single data frame as input for the visualization software.
- *Video collapsing* - Collapses video events in fixed intervals from the ‘DateTime’ values, and performs various aggregations and calculations for each group.
- *Obtaining sampling effort* - Processes video files from parent directories, calculates the duration and number of files for each subdirectory, organizes the results into a data frame, manipulates the data frame to extract relevant information, and writes the processed data to a .csv file for further merging with the video-level dataset.
- *Obtaining file metadata* - this code reads and processes video files’ metadata (e.g., video duration and creation time) from the parent directory, to merge the resulting metadata to the video-level dataset (e.g., EXIF tools).

#### Validation method

To assess the effectiveness of AI and evaluate data loss in its application, we used the recently compiled Frugivory-camtrap database, which offers a comprehensive overview of frugivory interactions within Doñana National Park (Villalva et al. 2024). This database was constructed using the AI workflow outlined in this paper. Due to the logistic and time limitations for visualizing the entire video-set in order to evaluate the performance of AI, we randomly selected 10% (*n*= 14,123 videos) of the raw files for an entire season (total sample size, *n_t_*= 138,351 videos), maintaining the species proportions, to assess binary classifications and build confusion matrices based on an established confidence threshold of 0.8. We computed fundamental performance metrics from the confusion matrices for each species (Table 2). Additionally, we incorporated performance indicators such as the F Score, widely used in statistical ecology (Chinchor 1992), and the MCC, recognized for its implementation in unbalanced datasets (Chicco & Jurman 2020). Confusion matrices were generated for each focal species and individual, allowing us to examine the information loss by analyzing false negatives and identifying potential reasons for detection failures.

**Table 2.** Common statistical rates and performance metrics to evaluate binary classifications through their confusion matrices. TP True positive, TN True negative, FP False Positive, FN False negative.

| Acronym | Description | Equation |
| --- | --- | --- |
| TPR | True positive rate / Sensitivity (Recall) | $TP/(TP+FN)$ |
| TNR | True negative rate / Specificity | $TN/(FP+TN)$ |
| FPR | False positive rate / Fall out | $FP/(FP+TN)$ |
| FNR | False negative rate / Miss rate | $FN/(FN+TP)$ |
| PPV | Positive predictive value / Precision | $TP/(TP+FP)$ |
| NPV | Negative predictive value | $TN/(TN+FN)$ |
| FDR | False discovery rate | $FP/(FP+TP)$ |
| FOR | False omisión rate | $FN/(FN+TN)$ |
| ACC | Accuracy | $(TP+TN)/(TP+TN+FP+FN)$ |
| ERR | Error rate | $(FP+FN)/(TP+TN+FP+FN)$ |
| F1 Score | Harmonic mean between TPR and PPV | $(2*TP)/((2*TP)+FP+FN)$ |
| MCC | Mathews correlation coefficient | $\frac{((TP*TN)-(FP*FN))}{\sqrt{((TP+FP)*(TP+FN)*(TN+FP)*TN+FN)}}$ |

In order to find the confidence threshold that optimizes the results in terms of reducing False positives while minimizing False negatives, we constructed a graph that combines success, True positive and False negative curves illustrating the optimal balance between the rate of success and the loss of information. Finally, we built the ecological networks with and without the inclusion of the missing data due to false negatives, providing insights into the extent and significance of data loss at both community and individual levels.

## 3. RESULTS

The primary output resulting from this article is the streamline protocol itself, which establishes a validated workflow facilitating the integration of computer vision to video recordings, addressing several crucial aspects such as data structuring and the incorporation of sampling effort, which are elaborated upon in the methods section (Fig. 1), as well as the accessory code useful for implementing the workflow that can be used as a tool in the GitHub repository (https://github.com/PJordano-Lab/Ecological-interactions-camtrap-protocol).

Upon visualizing and analyzing the randomly-stratified sample subset, the overall confusion matrix was obtained (see Table 2 for a conceptual summary of the confusion metrics). A confidence threshold of 0.80 resulted in the loss of 767 video clips containing animals. Out of this total, 81 clips were indeterminable due to various factors rendering them invisible in the footage (e.g., unknown species), not directly affecting our objective of gathering data for ecological interactions. Notably, 48% of the missing clips (*n* = 367) depicted various foraging behaviors (such as feeding, probable feeding, and searching for food), therefore representing legitimate missing data that directly influence data gathering for our purpose. Specifically, 83 clips depicted animals clearly engaged in fruit handling and feeding behaviors, while half of these animals were in the process of searching for fruit. The overall confusion matrix for the model (confidence level > 0.8) showed an accuracy of 0.92. However, there was significant variability in accuracy across different plant species ranging from 0.85 to 0.97 (Table 3).

**Table 3.** Confusion matrices, species-specific and combined, displayed in tabular format for a designated confidence threshold (c = 0.80) to depict how the AI model’s performance (figures refer to proportion of cases-no. of videos-falling in each category indicated as column names) varies across various species. no. videos refers to the size of a stratified random sample of the dataset that were visually revised (*n=* 14123), without AI assistance; such a sample was extracted from the total of *n*= 138,351 videos obtained. \**J. macrocarpa var. oxycedrus*.

| Plant species | True positive | True negative | False positive | False negative | no. videos |
| --- | --- | --- | --- | --- | --- |
| *Arbutus unedo* | 0.09 | 0.80 | 0.00 | 0.10 | 1431 |
| *Asparagus aphyllus* | 0.14 | 0.73 | 0.06 | 0.07 | 525 |
| *Corema album* | 0.05 | 0.92 | 0.01 | 0.02 | 2087 |
| *Juniperus macrocarpa* * | 0.08 | 0.87 | 0.01 | 0.04 | 391 |
| *Myrtus communis* | 0.01 | 0.96 | 0.00 | 0.03 | 936 |
| *Olea europaea* | 0.31 | 0.57 | 0.01 | 0.12 | 729 |
| *Pinus pinea* | 0.75 | 0.10 | 0.02 | 0.13 | 534 |
| *Pyrus bourgaeana* | 0.17 | 0.75 | 0.03 | 0.05 | 2670 |
| *Rubus ulmifolius* | 0.03 | 0.89 | 0.02 | 0.07 | 3357 |
| *Smilax aspera* | 0.01 | 0.95 | 0.02 | 0.02 | 1463 |
| Combined (all species) | 0.11 | 0.8 | 0.02 | 0.06 | 14123 |

The degree of missing information varied among species, with pairwise interactions declining by 1 to 10%. For example, in two distinct plant species-*Arbutus unedo* and *Pyrus bourgaeana-* the model’s failure also resulted in the loss of three unique interactions (Table 4). Correspondingly, performance metrics exhibited variability; certain species (e.g., *Olea europaea* and *Pyrus bourgaeana*) demonstrated high performance and consistent computer vision behavior, whereas others (e.g., *Smilax aspera*) performed poorly.

**Table 4.** Outcomes derived from video recordings capturing ecological interactions assisted by AI (RAW data) and the subsequent validation of 10% of the dataset (information lost). Results are grouped by focal plant species (rows) in three sets of columns detailing the (i) Total number of interaction events recorded, as supported by AI (excluding absent interactions), (ii) loss of information concerning pairwise interactions (number of interactions and species lost) and (iii) the loss of unique interactions (number of unique interactions and abbreviated name of species lost). Unique species lost: *Pica pica; Sturnus unicolor, Curruca undata; Sylvia atricapilla*. \**Curruca sp.* could belong to any species within the *Curruca* genus, yet it was not possible to discern the specific species from the video clip. The RAW dataset was constructed specifically for fleshy-fruited species, which is why *Pinus pinea* was excluded from this table.

| Plant Species | Raw data<br>(network) |  | Lost information<br>(Pairwise Interactions) |  | Lost information<br>(Unique interactions) |  |
| --- | --- | --- | --- | --- | --- | --- |
|  | No.<br>Inter<br>acti<br>ons | No.<br>Species | Interactions lost | Species<br>lost | Interactions<br>lost | Species<br>lost |
| *Arbutus unedo* | 222<br>9 | 24 | 64 (2.9 %) | 12 | 2 | *P. pica* (1)<br>*S. unicolor* (1) |
| *Asparagus aphyllus* | 777 | 13 | 16 (2.1 %) | 4 | 0 | - |
| *Corema album* | 656 | 18 | 13 (2.0 %) | 7 | 2 | *Curruca sp\** (1)<br>*C. undata* (1) |
| *Juniperus macrocarpa* | 270 | 9 | 3 (1.1 %) | 2 | 0 | - |
| *Myrtus communis* | 257 | 12 | 12 (4.7 %) | 6 | 0 | - |
| *Olea europaea* | 973 | 20 | 30 (3.1%) | 7 | 0 | - |
| *Pyrus bourgaeana* | 806 | 16 | 75 (9.3 %) | 11 | 2 | *S. atricapilla* (1)<br>*Curruca sp\** (1) |
| *Rubus ulmifolius* | 255<br>3 | 37 | 116 (4.5 %) | 14 | 0 | - |
| *Smilax aspera* | 150 | 11 | 15 (10.0 %) | 4 | 0 | - |

We were unable to discern the cause of failure in 29% of cases. However, we were able to determine that the majority (38%) of undetected species were obscured by vegetation in the background of the scene, likely eluding the AI’s detection. Interestingly, approximately 12% of species were overlooked by the AI probably due to their mimetic characteristics, such as the case of *Carduelis carduelis* in *Arbutus unedo*, where a substantial loss of interactions (90%) occurred likely because its colors resembled the prevalent yellow-red tones in *Arbutus unedo*. A total of three unique pairwise interactions were lost when using this protocol, namely *Arbutus unedo-Pica pica*, *Arbutus unedo-Sturnus unicolor* and *Pyrus bourgaeana*-*Sylvia atricapilla*. All three interactions involving animals in search for food were missed because the animal was located in the background of the scene.

We carried a direct comparison of the structure of the two networks resulting from the visualization protocol: one including the data with direct records of fruit handling/feeding and active fruit foraging (*n_t_=* 10,659 videos), and another including all the information, by imputing the potential false negatives (*FN*) (*n_fn_=* 11,574 videos). Both networks were described as adjacency matrices ***M****_t_* and ***M****_fn_* and their respective graphs *g_t_*, *g_fn_* were compared by quadratic assignment procedures (QAP; R package *sna*, function *qaptest*; Butts 2024). Function *qaptest* performs a repeatedly (randomly) relabeling the input graphs, recalculating the test statistic, and then evaluating the fraction of draws greater than or equal to (and less than or equal to) the observed value. We used two statistics for the comparisons, the Hamming distance (*H_d_*) and the graph correlation. Hamming distance is just the number of links that have to be moved in one graph to get the exact structure/topology of the other one. The graph correlation is useful to get a standardized measure of association between graphs (Butts and Carley 2005). For both statistics we obtained no evidence that the subsampled networks (without *FN* imputation) would result in structurally different network topologies than the networks including imputed *FN*s: *H_d_*= 831, *P<<* 0.0001, significantly lower than expected for different networks; and *gcor*= 0.9817, *P<<* 0.0001, suggesting a marked graph correlation between the two networks. Therefore, even ignoring *FN*s we could still accurately represent the structural properties of these complex networks; in other words, imputing *FN*s does not appear to overestimate or otherwise alter the link distribution and overall network structure of actual fruit-handling records.

## 4. DISCUSSION

Our protocol aims to obtain datasets tailored for the analysis of plant-animal interactions field records. But it is also applicable to other animal-focused studies, where video recording is required to capture details of animal behavior in natural environments. The application of computer vision to classify videos with/without animals enables the management of the vast amount of information generated in such approaches. The data structuring and processing proposed in this paper aid to minimize the effort/time required to obtain it. Through the use of this streamlined workflow, the time required to acquire a final database ready to analyze was notably reduced. In a pilot evaluation, we managed to decrease the time spent on viewing and annotating interaction videos from three weeks (part-time) to three days, thereby reducing the effort by 1/7, while achieving robust interaction data with minimal variations. Nevertheless, constructing databases with AI is not infallible, resulting in some information loss (i.e., loss of certain interactions because of undetected animal species by the computer vision model). Our results suggest that a significant portion of the information lost due to the application of AI does not contain relevant information for the objective of generating a robust ecological network database (species that were not identified in the scene by experts or species that presented behaviors different to active foraging for fruits). However, some of the information that contained relevant data (i.e., species exhibiting foraging behaviors that were not captured by AI) was certainly lost, posing a potential source of bias in the data for subsequent analysis. Importantly, the loss of information was not uniform across species. At the community level, some plant species exhibited more interaction losses than others and at the species level some plant individuals lost more interactions than others. This uneven loss of information potentially introduced bias into the obtained dataset, which may generate artifacts in subsequent analyses. For example, individual plants growing in specific conditions of vegetation cover, location of the light and shadows relative to the camera location, actual visibility of the individual fruits in the canopy, etc. The best procedure to avoid those potential biases would be replication of camera trap locations so that the full range of environmental and habitat conditions where plants are growing is captured. Undetected species lead to undetected interactions that might thus appear less central than they actually are in some plant individuals or even plant species, potentially misrepresenting their ecological role. This could potentially affect conclusions about species importance and interaction strength leading to estimation biases for network stability, resilience, and connectivity. Adequate replication and sufficient sampling effort (e.g., encompassing the whole fruiting phenophase) are the best ways to minimize such types of biases. In addition, care should be taken when deploying the cameras, in order to avoid potential artifacts affecting detection (e.g., obtrusive branches and leaves, sunflecks, excessive shade, etc.). Even though these results were obtained using the MegaDetector model, it is likely that similar context-dependent patterns will appear when using other object detection models. This highlights the importance of validating AI models in specific ecological contexts to ensure data accuracy.

The loss of information caused by AI can not easily be eradicated. In many cases, animal species were present in the scene, but were located in the background and only recognizable by movement patterns, size, sound, or by context dependent and species-specific characteristics that triggered open discussions among expert ornithologists. These contingencies are difficult to emulate through computer vision models and will be unrealistic trying to avoid them regardless of how detailed the training dataset is. In some cases, the loss of data was attributed to the mimicry of certain animal species on specific plant species. A good example occurs with *Arbutus unedo* and *Carduelis carduelis*, where facial and wing colors (red, cream, black, and yellow) closely resemble the reddish-orange tones of the scrub fruits, resulting in a 90% loss of information for this particular interaction. This phenomenon, clearly evident in this species, may be happening among mimetic species such as certain thrush species (*Turdus philomelos* or *Turdus torquatus*), which may go unnoticed especially when standing in the bare ground compared to *Turdus merula*, generally much more visible, potentially biasing the database. In any case, in our experience, those instances occurred infrequently and, again, replication and sufficient sampling effort may counteract potential biases.

The approach outlined in this study proves particularly usable for constructing community-level, species interaction networks, as the extent of information loss was minimal in comparison to the substantial benefit gained from acquiring extensive data. While some information was lost (e.g., variations in interaction frequencies), the sheer volume of data generated diluted the loss of interactions, minimizing its significance. Previous analyses of sample coverage using these camera-trap based methods (Villalva et al. 2024, Quintero et al. 2024, Isla et al. 2024) evidenced sufficient sampling and adequate estimation of the actual frequency of interactions. Our analysis regarding the validation of AI showed that the overall network structure is a faithful reflection and therefore offers homologous results, to non AI processing for our purposes (Figure 4). However, there remains a loss of unique interactions that could be important, especially from the zoocentric perspective, as this biological information about animal frugivory and fruit resource use will be lost. Nevertheless, from a phytocentric viewpoint and considering the entire frugivore community, these interactions may be less critical, given that the most common seed dispersers are those typically playing a predominant role in maintaining fleshy-fruited tree species’ persistence (Rehling et al., 2023).

**Figure 4.**
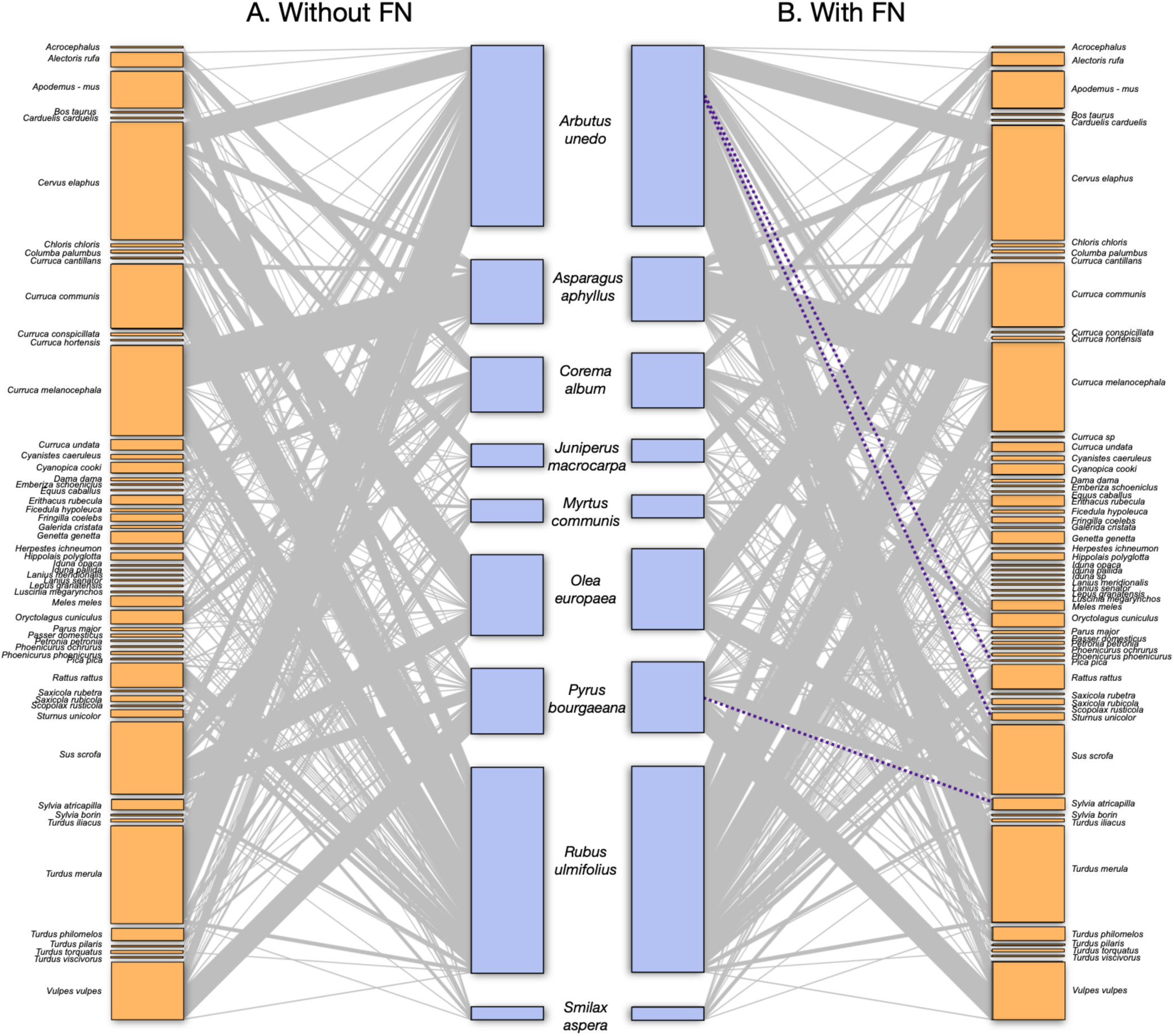
Community level network built using AI (left) and with the inclusion of False negatives (right). Unique pairwise interactions missed by applying AI are highlighted in purple dotted lines. The size of the rectangles representing plants (blue) and animals (orange) nodes reflects the frequency of events involving each individual or species respectively. The thickness of connections indicates the relative frequency of each pairwise interaction.

**Figure 5.**
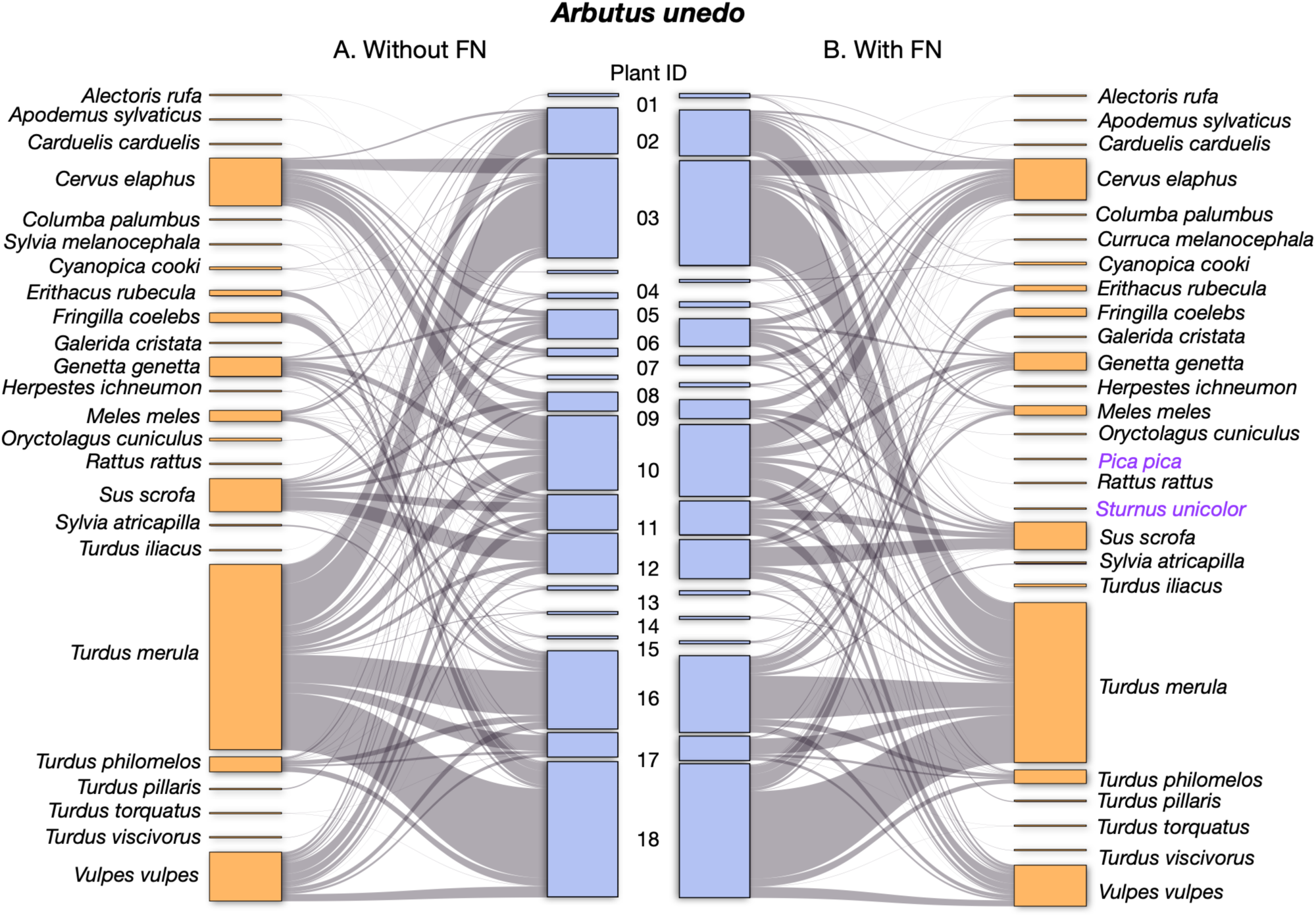
Individual-based network of plant-animal interactions including false negatives (right) contrasting with one constructed without false negatives (left). Individual plants of *Arbutus unedo* are represented as blue rectangles positioned in the center (numbers identify individual plants sampled). Animal species nodes are aligned at the margins, depicted as orange rectangles. The size of the rectangles reflects the frequency of events involving each plant individual or animal species. The thickness of connections (grey) indicates the relative frequency of each pairwise interaction. Animal species common to both networks are depicted in black, while those present in one network but not the other are highlighted in purple.

When constructing networks at the individual level (i.e., documenting frugivore species interacting with different plant individuals in a local population), certain limitations may arise. As stated above, information loss was not evenly distributed, being more pronounced in specific individuals. The impact of interaction loss on certain individuals can substantially influence analysis outcomes, especially in individual-based approaches, thereby introducing significant bias. To mitigate this effect, employing additional complementary methods alongside camera traps, such as direct observations, mist netting, or scat barcoding, can provide a more comprehensive inventory of individual assemblages. However, merging datasets from various methods can be intricate and presents limitations, particularly concerning quantifying interaction intensity in relation to the specific ecological questions addressed in each study (Quintero et al., 2022).

Our results also indicate that choosing a confidence threshold up to which to review is not a trivial choice, especially in cases where the large amount of videos makes it impossible to lower the confidence levels. Therefore, it is necessary to ensure the robustness of the data obtained in these cases. To achieve this, it is necessary to reduce the effort in viewing and tagging video sequences while maintaining data robustness. Thus, it is necessary to optimize visualization, reducing the false negative rate (information loss-sequences excluded that actually contain interaction records), which decreases with model confidence, and the false positive rate (empty videos lacking actual interactions but that we review and therefore represent a loss of time), which increases with model confidence. To achieve this, we advise to visualize a random sample of the video-set to obtain the confusion matrix and plot cumulative curves depicting false positive/negative rates relative to increasing confidence levels, allowing to find the cut-off point between both rates. This will represent the optimal value for setting a confidence threshold for optimization. Importantly, our results point out that this confidence level is different for each plant species, and may range between ca. 0.25-0.75. Setting the same level for all of them (a common practice when the use of AI is trivialized) would introduce a serious bias in the lost information for different focal species. Therefore, it is important to use a particular confidence level for different focal species, with the proposed a priori assessment of this threshold being determined from an adequately designed random sample of the available video recordings. It is likely that this variation results from different plant structure and background (Palencia et al. 2022) and that it also may occur in different scenarios or contexts, as has been highlighted in studies finding variations in the detection for different animal species regardless of the background (e.g., Vélez et al., 2023).

Most of the scientific literature on the application of AI to camera traps is focused on citing the accuracy of predicted values as the main predictive metric (Norouzzadeh et al. 2018, Schneider et al. 2020, Tabak et al. 2019). However, this measure is certainly not an adequate approximation to the quality of the data required for an approach like ours. Accuracy may be unbalanced, where some species will have far more errors than others (Greenberg, 2020), especially in the case of unbalanced databases where the number of cases of one class (usually negatives) is much greater than the rest of the cases. In such cases accuracy provides a very optimistic estimate of the classifier’s ability on the dominant class and is not an indicator of the model’s performance. The same overoptimistic effect occurs when using the F1 performance metric, which is not suitable for this type of unbalanced data. The MCC (Matthew Correlation Coefficient) can be more appropriate for unbalanced confusion matrices and has been already applied to binary classification for genomic scenarios (Chicco and Jurman, 2020). This metric produces a more informative and trustworthy score in evaluating binary classifications than accuracy and F1 score incorporating the class imbalance component into its mathematical properties, requiring correct classifications in both positive and negative cases, regardless of the ratios of the complete dataset.

Image recognition models continue to be enhanced with new data, leading to better performance over time. For instance, the newer version of MegaDetector (MDv5) increased processing speed and incorporated additional training data to improve detection of particular taxa (rodents, reptiles, and small birds). These updates, along with advancements in AI platforms for data management and integration, will further enhance the efficiency of ecological studies in the near future.

In the current context of rampant biodiversity loss, there is a strong interest in minimizing human impacts on biodiversity. Significant efforts are underway to curb this issue, as reflected in the 23 targets for preventing future biodiversity loss set in the 2022 Kunming-Montreal Global Biodiversity Framework (GBF). Substantial efforts have been made to identify and standardize indicators that reflect biodiversity changes. Although there are many Essential Biodiversity Variables (EBVs), most oversimplify biodiversity (as through just listing species or habitats trends) that do not fully capture the complexity of ecosystems nor their ecological functions, which is a crucial aspect for biodiversity conservation (Bullock et al., 2022). Therefore, presenting standardized methods that incorporate the latest technological advances—such as this protocol—holds great potential for broad-scale application (i.e. in national assessments of community structure). These methods offer highly relevant information for understanding and monitoring changes in community structure and biodiversity of ecological interactions and functions, thereby aiding in the understanding and quantification of biodiversity loss and with a special value for restoration of ecological processes. If implemented at broad-scale, next steps could also include the combination with remote sensing as a complement to understand changes in vegetal communities colonization fronts and their link to biodiversity complexity.

The protocol presented in this publication, designed to generate field databases for analyzing plant-animal ecological interactions networks, has proven to be an optimization for robust ecological interaction field data collection. It incorporates a clear and standardized workflow, building on prior work on camera trap standards and publicly available AI-powered platforms. It provides specific guidelines for setting up cameras in the field, organizing data to automate data extraction (i.e., sampling effort), and using computer vision to classify videos with and without animals as well as optimizing the visualization and tagging process. This approach significantly reduces the time and effort required to process and analyze field data, resulting in a final dataset ready for network analysis or other behavioral studies. Additionally, we offer methods to identify the confidence threshold for different focal species to balance false negatives and false positives, and we discuss the main caveats of using AI for this type of dataset, particularly the flaws that may arise in individual-based approaches, while highlighting the robustness for community-level approaches.

## ACKNOWLEDGEMENTS

We appreciate the help in the field and extensive discussions and ideas of Francisco Rodríguez-Sánchez, Eva Moracho, Blanca Arroyo-Correa, Miguel Jácome, Elena Quintero, Pablo Homet, Jorge Isla, Irene Mendoza, Gemma Calvo, and many others. Field work was greatly assisted by Gemma Calvo, Pablo Homet and Eugene W, Schupp. Additionally, we thank Ousamma Tourabi, Hammad Zubahir-Sheikh, and Margaret Hemp for their help in revising the stratified random sampled videos. We are in debt with agent Dan Morris that helped and discussed how to implement Medgatector for videos and kindly helped running large video batches. Over the years our field work has been generously supported by the facilities ICTS-Reserva Biológica de Doñana (CSIC) and the administration of the Doñana National Park. Completion of this manuscript was funded by grants PID2022-136812NB-I00 from the Agencia Estatal de Investigación, the Plan Propio de Investigación y Transferencia, University of Sevilla (2021-2024), and a LifeWatch ERIC-SUMHAL project (LIFEWATCH-2019-09-CSIC-13) with FEDER-EU530 funding and PV considers this work as a contribution to SustainScapes (grant NNF20OC0059595).

## Supplementary Materials

### S.1 Database structure

Most likely, a field sampling using camera traps for recording ecological interactions involves the simultaneous deployment of multiple cameras, in replicated positions to target different individual plants, throughout the fruiting season of different focal plant species. The cameras are checked at regular intervals, typically weekly, biweekly, or monthly, which can result in a large number of videos with the same name and date.

Effective management of such a large and complex data set is crucial for a successful database creation. To achieve this, a structured field database is required to keep track of the data at every stage of the process. For this purpose, we recommend the use of the Camera Trap Data Package (Camtrap DP), a community-developed data exchange format that is under development as a Biodiversity Information Standard (TDWG). This package provides a useful structure for controlling camera-trap data at three levels, from which we will adopt the structure: *deployments*, *revisions*, and *observations*.

Camtrap DP offers a standardized format for organizing camera-trap data, ensuring consistency and reducing the risk of errors or inconsistencies. This structure includes all necessary information, such as camera settings, deployment locations, and video file names, allowing for easy management and analysis of the data. A template for the Camtrap DP is available for use, and you can find a template for our ad-hoc structure in the GitHub repository (https://github.com/PJordano-Lab/Ecological-interactions-camtrap-protocol/tree/main/Preprocess or see the following descriptions.

The main data is structured in three related plain text files (.csv) as follows:

| File | Description |
| --- | --- |
| deployments.csv | Table with camera trap deployments. |
| video.csv | Table with media files captured by the camera traps. |
| observations.csv | Table with observations based on media files (after viewing in Timelapse) |

#### S.1.1 Deployments

Table with camera trap deployments. Includes deploymentID (focal species acronym + cameraID), Location and camera Setup information for each camera.

| Name | Definition | Type |
| --- | --- | --- |
| Deployment_ID | Unique identifier of the deployment. Name of the focal species followed by the individual number low dash camera number.<br>Example: Aune003_58 | string |
| Location | Name given to the deployment location. Survey area. | string |
| Longitude | Longitude of the deployment location in decimal degrees, using the WGS84 datum. | number |
| Latitude | Latitude of the deployment location in decimal degrees, using the WGS84 datum. | number |
| Start | Date and time at which the deployment was started. Formatted as an ISO 8601 string with timezone designator (YYYY-MM-DDThh:mm:ss±hh:mm). | datetime |
| End | Date and time at which the deployment was ended. Formatted as an ISO 8601 string with timezone designator (YYYY-MM-DDThh:mm:ss±hh:mm) | datetime |
| Days | Number of days the deployment was set in the field. End_date - Start_date | number |
| Setup_by | Name(s) or unique identifier of the person that deployed the camera | string |
| Camera_ID | Unique identifier of the camera used for the deployment (could be the serial number but also a simple number) | string |
| Camera_model | Manufacturer and model of the camera | string |
| Comments | Comments or notes about the deployment. | string |

#### S.1.2 Videos

Table with video files captured by camera traps. Associated with deployments (by deploymentID) and organised in revisions (revision_ID). Includes Timestamp_Issues and File_path.

| Name | Definition | Type |
| --- | --- | --- |
| Deployment_ID | Unique identifier of the deployment the media file belongs to. Foreign key to Deployments.Deployment_ID | string |
| Revision_ID | Unique identifier of the revision the media file belongs to. Revisions contain one or more media files (e.g. a single image or video or a sequence of successive images or videos). Example: Rev_01 | string |
| Videos | Number of media files | number |
| First_video | Datetime for the first video in the revision sequence | daytime |
| Last_video | Datetime for the last video in the revision sequence | daytime |
| Days | Number of days that the deployment was set in the field during the current revision | number |
| Setup_date | Date at which the camera was set on in the current revision. Date at which the camera was set off in the current revision | daytime |
| Revision_date | Date at which the camera was set off in the current revision. Date at which the camera was set off in the current revision | daytime |
| Functioning_days | Number of days where the camera was functioning. Revision_date - Setup_date | number |
| Battery | Percentage of battery in the revision datetime | string |
| Timestamp_Issues | True if timestamp in the media have been detected | boolean |
| File_path | URL or relative path to the media files, respectively for externally hosted files | string |
| Favourite | True if it contains videos tagged as favorite | boolean |
| Comments | Comments or notes about the revision | string |

#### S.1.3 Observations

Table with video files results from visualization. Associated with deployments (deploymentID) and with revisions (revision_ID) through Videos.file_path. Note that this data will be generated directly with Timelapse software as explained below.

| Name | Definition | Type |
| --- | --- | --- |
| File | Name of the video file. If more than one video files use concatenate separated by “,” | string |
| Path | URL or Relative path to the first **Obs.File**, respectively for externally hosted files | string |
| Plant_sp | Name of the focal plant species | string |
| Plant_ID | Unique identifier of the plant individual. Example: Sasp003 | string |
| DateTime | Date and time at which the video started. Formatted as an ISO 8601 string with timezone designator (YYYY-MM-DDThh:mm:ss±hh:mm) | datetime |
| Sp1 | Latin binomial for the principal animal species recorded in the video. Example: Athene noctua | string |
| Behaviour |  | string |
| Sp2 | Latin binomial for a secondary animal species recorded in the video | string |
| Behaviour_Sp2 |  | string |
| Sp3 | Latin binomial for a secondary animal species recorded in the video | string |
| Behaviour_Sp3 |  | string |
| N_cam | Number of cameras set in the same focus individual | number |
| Videos | Number of videos recorded in the current revision | number |
| Days | Number of days for the camera recording in the current revision. Sampling effort for a video in a given revision | number |
| Video_events | When videos are collapsed in time (i.e 5 minutes) number of videos that where collapsed. It will be coincident with the number of different **Obs.File** names | number |
| Duration | Number of second for the duration of the event. Could be the sum of durations from different **Obs.video_events** | number |
| TimeStamp_Issue | True if timestamp in the video have been detected |  |
| Long | Longitude of the deployment location in decimal degrees, using the WGS84 datum | number |
| Lat | Latitude of the deployment location in decimal degrees, using the WGS84 datum | number |

## Notes

https://github.com/PJordano-Lab

### Competing Interest Statement

The authors have declared no competing interest.

